# Connect-seq to superimpose molecular on anatomical neural circuit maps

**DOI:** 10.1101/454835

**Authors:** Naresh K. Hanchate, Eun Jeong Lee, Andria Ellis, Kunio Kondoh, Donghui Kuang, Ryan Basom, Cole Trapnell, Linda B. Buck

## Abstract

The mouse brain contains ~100 million neurons interconnected in a vast array of neural circuits. The identities and functions of individual neuronal components of most circuits are undefined. Here we describe a method, termed ‘Connect-seq’, which combines retrograde viral tracing and single cell transcriptomics to uncover the molecular identities of upstream neurons in a specific circuit and the signaling molecules they use to communicate. Connect-seq can generate a molecular map that can be superimposed on a neuroanatomical map to permit molecular and genetic interrogation of how the neuronal components of a circuit control its function. Application of this method to hypothalamic neurons controlling physiological responses to fear and stress reveal subsets of upstream neurons that express diverse constellations of signaling molecules and can be distinguished by their anatomical locations.

## Introduction

The brain contains a multitude of neural circuits that control diverse functions, but the specific neuronal components of most circuits and their respective roles are largely a mystery^1^^-^^3^. The application of single cell RNA sequencing techniques has allowed rapid advances in defining the transcriptomes of brain neurons^1^^-^^6^. However, much less is known about which of the vast numbers of neurons with known transcriptomes are interconnected to control specific functions. Here we devised a strategy, “Connect-seq”, to obtain additional information about the neuronal components of specified neural circuits. The strategy couples the use of conditional neurotropic viruses that cross synapses with transcriptome analysis of connected upstream neurons infected with virus to obtain the molecular identities of neurons connected in a circuit.

To test the strategy, we focused on neural circuits that transmit signals to corticotropin releasing hormone neurons (CRHNs) in the paraventricular nucleus of the hypothalamus (PVN)^7^. In response to danger, CRHNs induce increases in blood stress hormones that act on multiple tissue systems to coordinate appropriate physiological responses. Surges in blood stress hormones occur in response to a variety of external and internal dangers or ‘stressors’, including predator odors, tissue injury, inflammation, and emotional and psychological stress^7^^,^^8^. Studies using classical neuroanatomical and neurophysiological approaches have provided a large body of information on PVN inputs that may affect CRHNs and the effects of classical neurotransmitters on CRHN activity and function^7^^,^^9^^-^^12^. However, the exact identities of upstream neurons that control CRHN functions and the mechanisms by which they do so have not been defined.

Viral tracing studies indicate that CRHNs receive direct synaptic input from neurons in 31 different brain areas^8^. These include 19 areas of the hypothalamus, a brain area that integrates information from the body and external environment and organizes fitting behavioral and physiological responses. Neurons two or more synapses upstream of CRHNs are seen in additional areas, including one small area of the olfactory cortex that has proved key to stress hormone responses to predator odors^8^. The viral tracing studies provide an anatomical map of neurons directly presynaptic to CRHNs, but the molecular identities of those neurons are unknown.

If one could identify genes whose expression defines subsets of neurons upstream of CRHNs in different brain areas, one would have molecular tools to explore which subsets mediate responses to different stressors and to manipulate those subsets to gain insight into how they function. Information regarding neurotransmitters and neuromodulators used by the upstream neurons to communicate with CRHNs would, in addition, provide information relevant not only to an understanding of neural circuit functions, but also potential insights into pharmacological interventions to modify the functions of specific circuit components.

To test the Connect-seq strategy, we infected Cre recombinase-expressing CRHNs with a Cre-dependent Pseudorabies virus (PRV) that travels retrogradely across synapses. Using flow cytometry, we isolated single virus-infected neurons and then used single cell RNA sequencing (RNA-seq) to define the transcriptomes of individual upstream neurons. These experiments revealed an enormous diversity of neurotransmitter and neuromodulator signaling molecules in neurons directly upstream of CRHNs, including more than 40 different neuropeptides. Many individual neurons coexpressed multiple different signaling molecules. These included neurons coexpressing neuropeptides with neurotransmitters or biogenic amines, as well as neurons coexpressing different neuropeptides. The combinations of signaling molecules expressed by different cells were extremely diverse. Upstream neurons expressing specific signaling molecules mapped to selected brain areas, demonstrating that Connect-seq can provide molecular tools with which to dissect the functions of individual neuronal components of neural circuits.

## RESULTS

### Transcriptome analysis of neurons upstream of CRHNs

We first developed a Pseudorabies virus, PRVB180, which has Cre recombinase-dependent expression of thymidine kinase (TK) fused to green fluorescent protein (GFP) (Fig. 1a). After infecting neurons expressing Cre, this virus will travel retrogradely across synapses to infect upstream neurons. We injected PRVB180 into the PVN of CRH-IRES-Cre (CRH-Cre) mice (Fig. 1b), which express Cre in CRHNs^13^. On day 3 post infection (d3pi), immunostaining of brain sections for GFP showed GFP+ (PRV+) neurons in the same brain areas as seen previously for PRVB177, which expresses TK-HA instead of TK-GFP and travels to directly presynaptic neurons on d3pi^8^.

**Figure 1.**
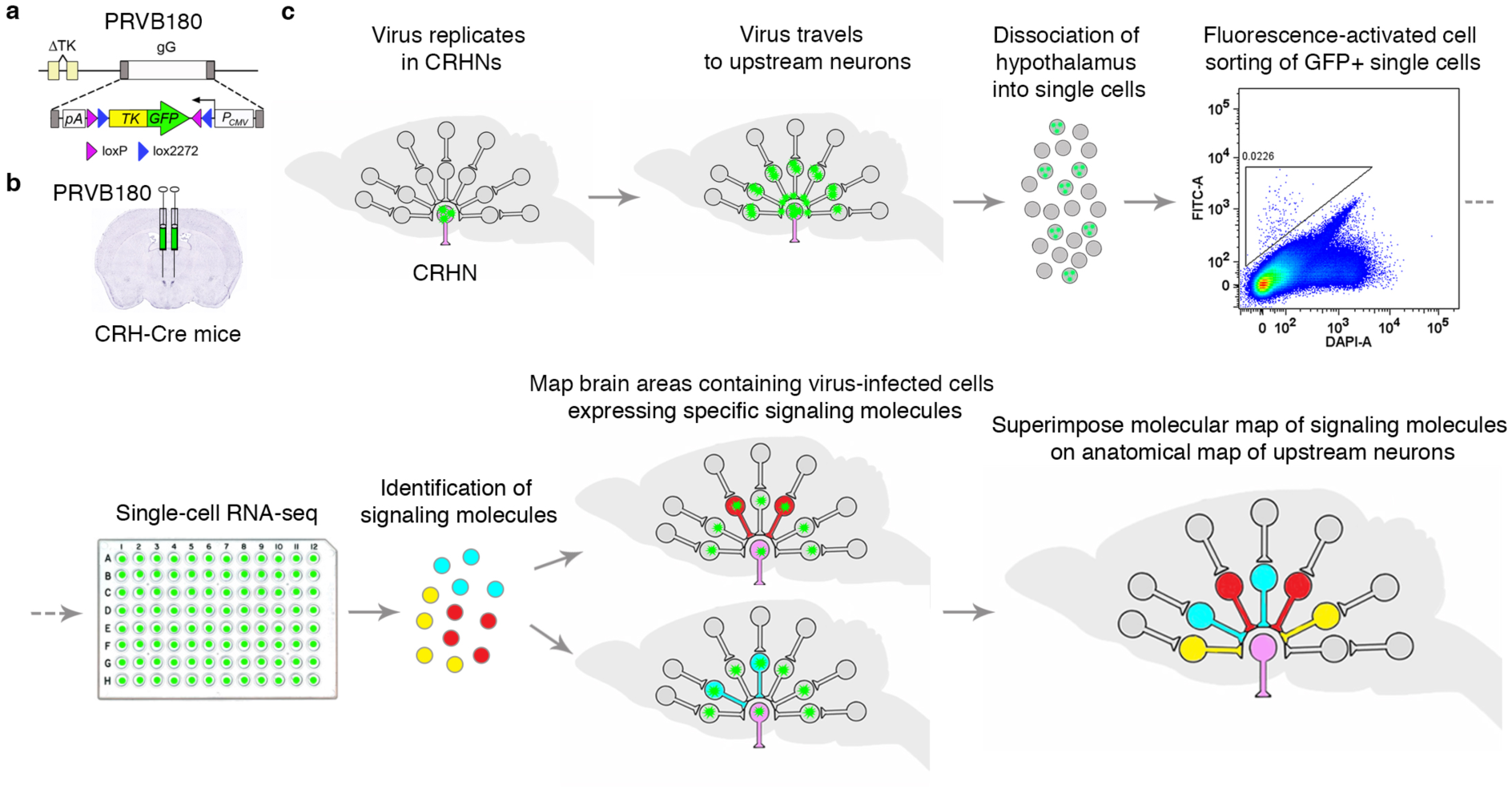
Connect-seq method. **(a)** PRVB180 has Cre recombinase-dependent expression of a thymidine kinase-green fluorescent protein fusion protein (TK-GFP). **(b)** CRHNs were infected with PRVB180 by injecting the virus into the PVN of CRH-Cre mice. **(c)** Following PRVB180 replication in CRHNs and travel of virus to upstream neurons, the hypothalamus was isolated and dissociated to obtain single cells. Flow cytometry was used to deposit cells with high GFP (FITC-A) and low DAPI-A staining, one per well, in multiwell plates. RNA-seq was used to determine their molecular identities and the signaling molecules (neurotransmitters/neuromodulators) they expressed. The locations of upstream neurons expressing selected signaling molecules were then determined, allowing a molecular map of upstream neurons to be superimposed on the existing anatomical circuit map.

We next conducted transcriptome analysis of neurons upstream of CRHNs. We infected CRHNs with PRVB180 and on d3pi isolated the hypothalamus and dissociated the tissue into a single cell suspension (Fig. 1c). Using flow cytometry, we isolated GFP+ (PRV-infected) cells, one per well, in 96 well plates^14^. We then conducted RNA-seq on each cell by Illumina sequencing cDNAs prepared from each cell^15^^,^^16^. Gene expression in individual cells was determined by alignment to the mouse genome using standard methods^17^^,^^18^. We proceeded to analyze 124 cells that expressed neuronal markers but not markers of other cell types, such as glial cells (see Methods).

### Neurons upstream of CRHNs express diverse neurotransmitters and neuromodulators

To characterize the neurons upstream of CRHNs, we focused on signaling molecules that neurons use to convey information to their downstream partners^19^. These include classical ‘fast neurotransmitters’, which act via ligand-gated ion channels on downstream neurons to rapidly activate or inhibit those neurons, and numerous neuromodulators that bind to G protein-coupled receptors (GPCRs) on downstream neurons to modulate their excitability. We reasoned that the large number of different signaling molecules in the brain and their differential expression among neurons could serve to 1) optimize the discovery of molecular identifiers that would distinguish different upstream neurons and 2) provide potential insight into the molecular mechanisms used by upstream neurons to communicate with CRHNs.

We identified a large number and variety of neurotransmitters and neuromodulators expressed by PRV-infected neurons upstream of CRHNs (Fig. 2). These included glutamate and GABA, which act via ligand-gated ion channels and are the major excitatory and inhibitory neurotransmitters in the brain, respectively. We also identified two other fast neurotransmitters that signal through ligand-gated ion channels, acetylcholine and glycine (Fig. 2a, 2b). All of these neurotransmitters can also act as neuromodulators by binding to specific GPCRs on downstream neurons.

**Figure 2.**
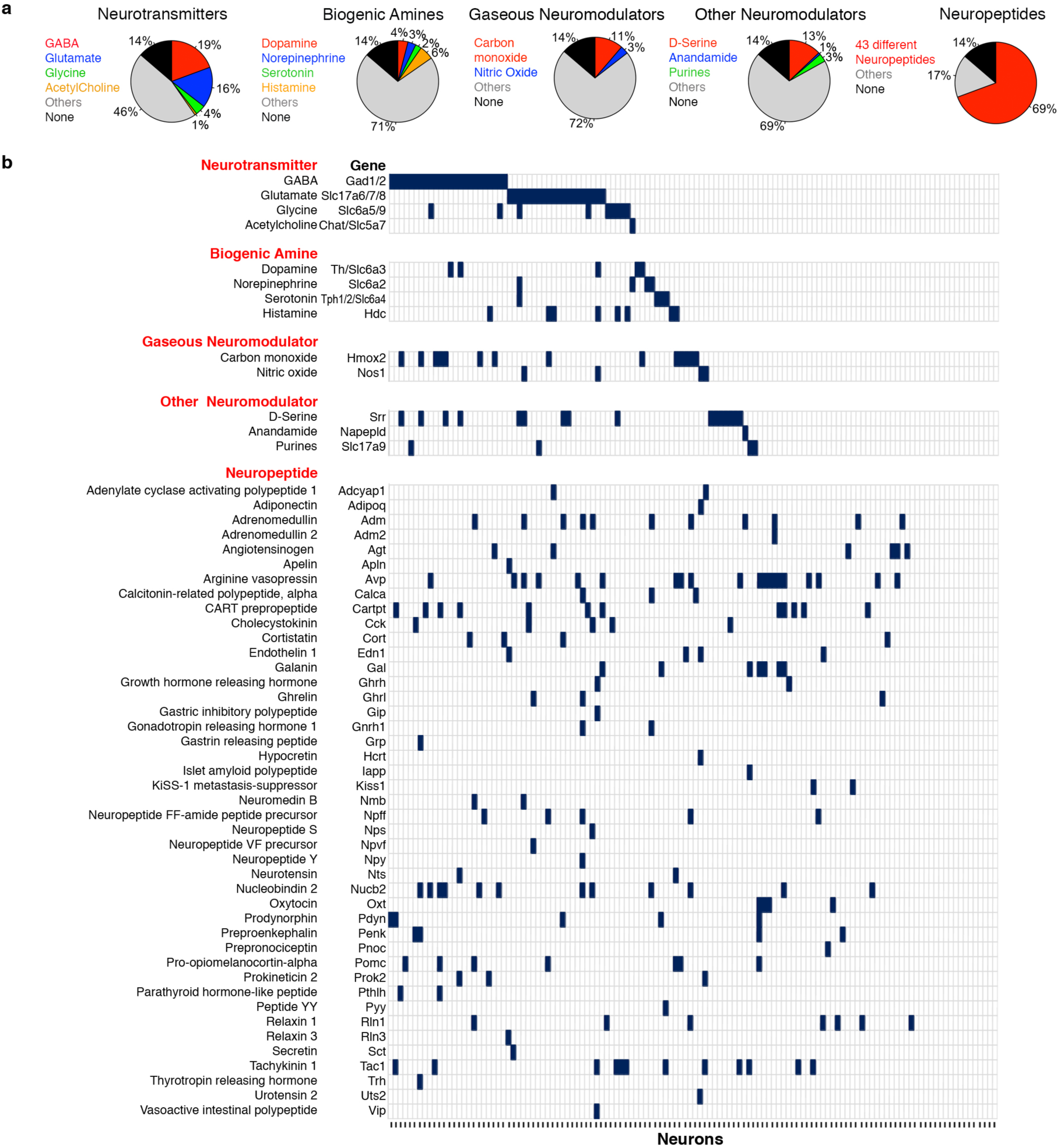
Neurons upstream of CRHNs express a large array of signaling molecules. **(a)** Pie charts show the percentages of upstream neurons expressing different neurotransmitters, biogenic amines, gaseous neuromodulators, other neuromodulators, and neuropeptides. **(b)** Signaling molecules expressed in 124 individual upstream neurons. Different neurons are indicated on the x-axis and different signaling molecules and their encoding or indicator genes on the y-axis. Vertical bars indicate expression at ≥10 FPKM (fragments per kilobase of transcript per million mapped reads).

The neurotransmitters were expressed in 40.3% of upstream neurons analyzed (Fig. 2a), but the number of neurons expressing different neurotransmitters varied (Fig. 3a). Glutamate and GABA were expressed in 16.1% and 19.4% of the neurons, respectively, while the other neurotransmitters were expressed in far fewer neurons (0.8% for acetylcholine and 4.0% for glycine) (Fig. 2a). CRHNs have ligand-gated channels for both glutamate^20^ and GABA^21^^,^^22^. And receive direct synaptic input from both neurotransmitters^11^^,^^23^^,^^24^. Our results suggest that CRHNs can be activated by many glutamatergic presynaptic neurons, but that many other upstream neurons may suppress CRHN activity via GABA transmission. Previous studies indicate that upstream GABAergic neurons can provide tonic inhibition to CRHNs, the release of which can lead to CRHN activation by disinhibition^11^^,^^25^. In addition, some upstream GABAergic neurons might be involved in stimulus-induced suppression of CRHN responses, as previously reported for certain odors^26^^,^^27^.

**Figure 3.**
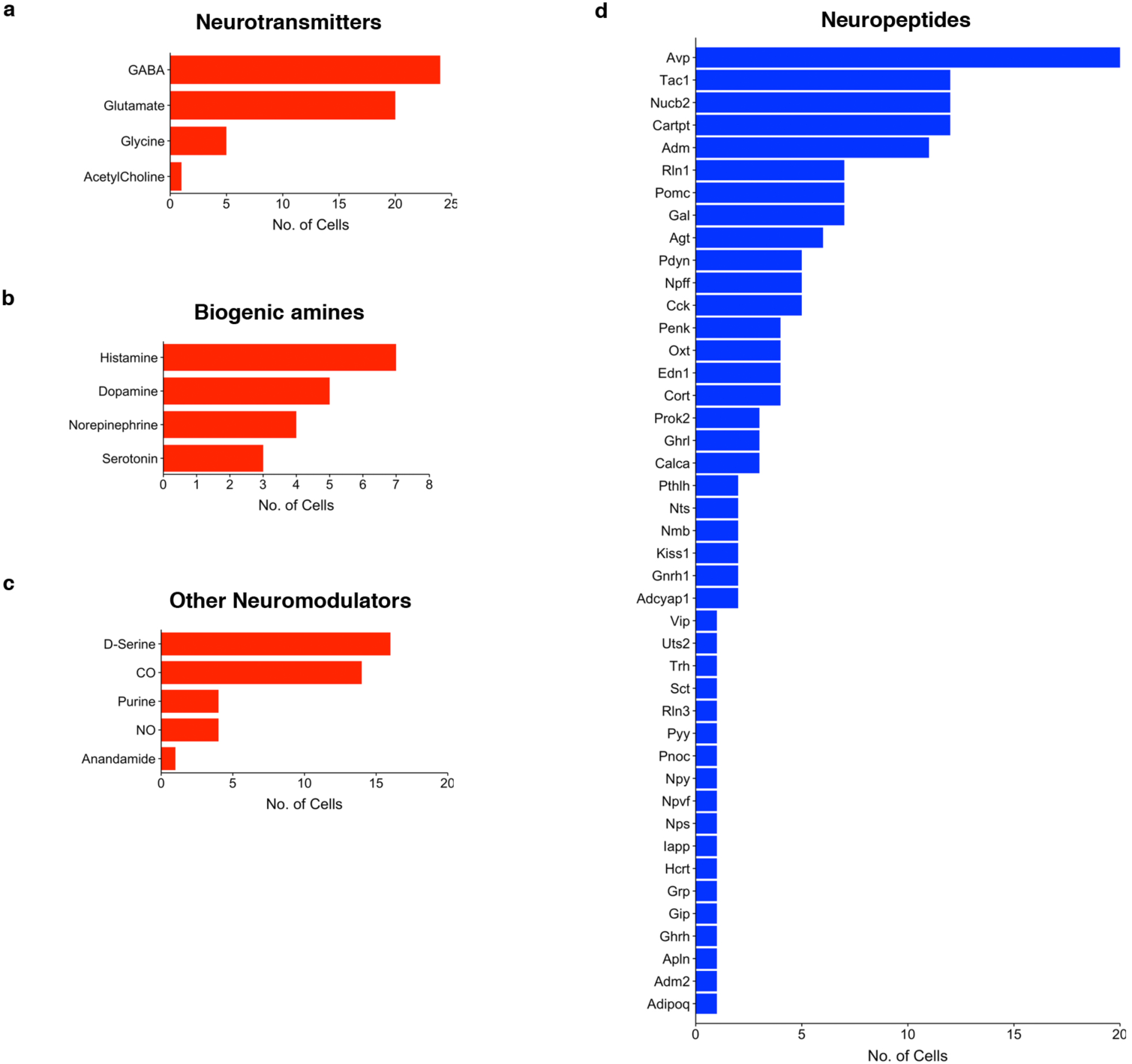
Different signaling molecules are expressed in different proportions of upstream neurons. **(a-d)** Graphs show the number of upstream neurons expressing different neurotransmitters (a), biogenic amines (b), other neuromodulators (c), and neuropeptides (d).

Small numbers of upstream neurons expressed biogenic amines (dopamine, norepinephrine, serotonin, or histamine) (Fig. 2b, 3b). These molecules, which signal through GPCRs, have widespread effects in the brain, can also act on peripheral tissues, and are the targets of numerous pharmacological agents^28^. Interestingly, several biogenic amines have reported effects on stress hormones^7^^,^^11^ and previous studies indicate that CRHNs have serotonin receptors and can be depolarized by serotonin^29^. By identifying the functions of different biogenic amine-expressing neurons upstream of CRHNs and determining how the amines affect CRHNs, it is possible that small molecule agonists or antagonists for specific biogenic amine receptors or their transporters could ultimately be employed to block or enhance the effects of particular stimuli on CRHNs.

Less common types of neurotransmitters/neuromodulators were also identified in small numbers of upstream neurons. These included the gaseous neuromodulators carbon monoxide and nitric oxide, as well as D-serine, the cannabinoid anandamide, and the purines ATP and adenosine (Fig. 2b, 3c).

Remarkably, 43 different neuropeptides were identified in neurons upstream of CRHNs (Fig. 2b, 3d). Neuropeptides can act as neuromodulators to enhance or dampen neuronal excitability by binding to GPCRs on downstream neurons ^30^^,^^31^. Neuropeptide expression was seen in 69.4% of upstream neurons (Fig. 2a). The number of neurons expressing different neuropeptides ranged from a single neuron (e.g. gastrin releasing peptide and neuropeptide Y) to 20 neurons (arginine vasopressin) (Fig. 2b, 3d). The large number of neuropeptides identified in upstream neurons and their differential expression among the upstream neurons provides a rich set of tools for superimposing a molecular map on the anatomical map of CRHN upstream circuits and for dissecting the functions of different subsets of upstream neurons.

### Upstream neurons can express multiple signaling molecules

Many of the upstream neurons expressed more than one neurotransmitter and/or neuromodulator (Fig. 4a and Supplementary Fig.1). More than 85% of neurons expressed at least one of the signaling molecules examined (Fig. 4b). Strikingly, 55% of the neurons expressed two or more signaling molecules and 21.8% expressed four or more (Fig. 2b, 4b).

**Figure 4.**
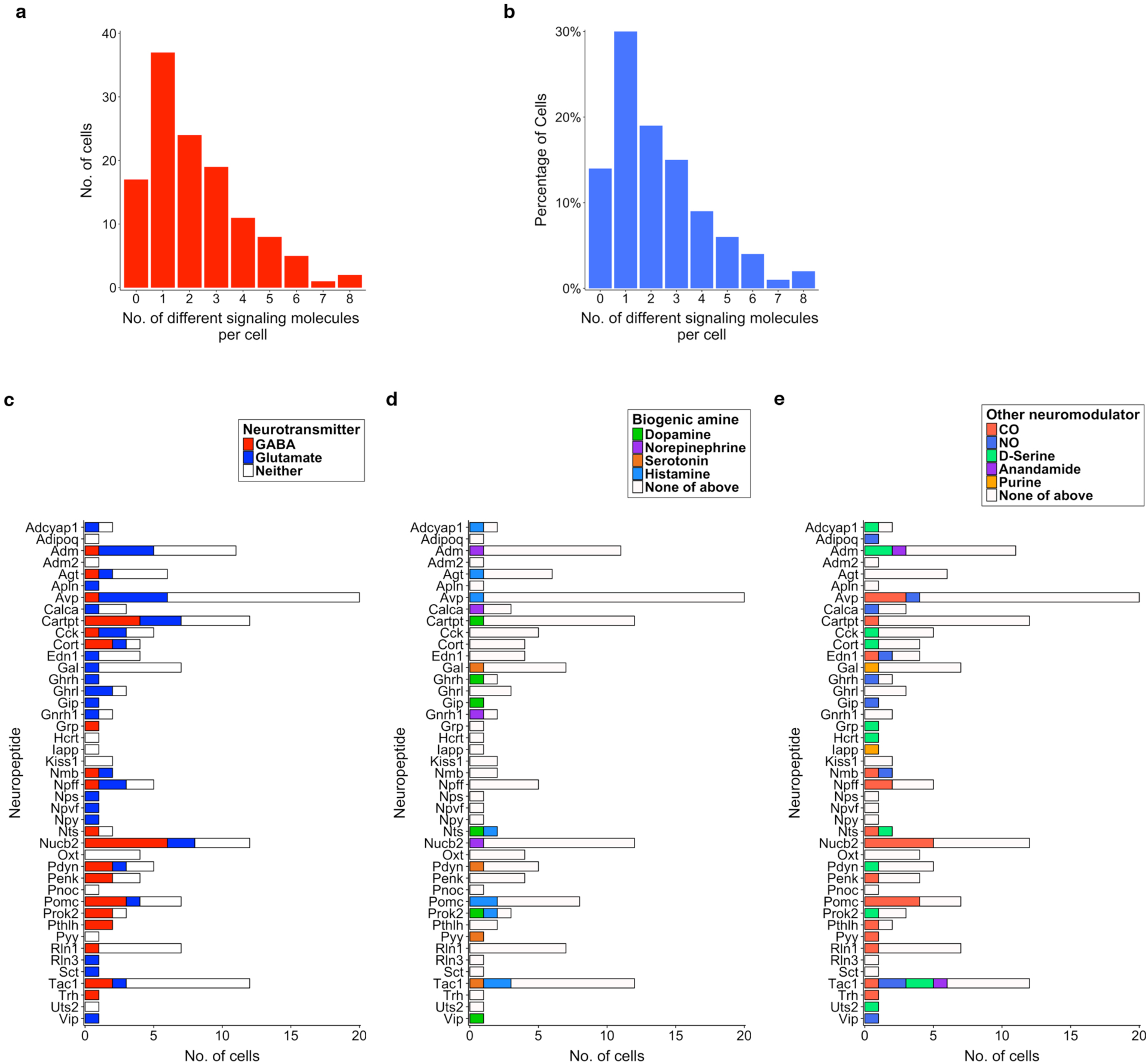
Combinatorial expression of signaling molecules in neurons upstream of CRHNs. **(a-b)** Graphs show the number (a) and percentage (b) of upstream neurons expressing different numbers of neurotransmitters and/or neuromodulators. **(c-e)** Graphs show the number of cells that coexpressed different neuropeptides with the excitatory neurotransmitter, glutamate or the inhibitory neurotransmitter, GABA (c) or with different neuromodulatory biogenic amines (d) or other neuromodulators (e).

Many upstream neurons coexpressed a given neuropeptide together with glutamate or GABA (Fig. 2b, 4c) and/or, in some cases, a biogenic amine (Fig. 2b, 4d) or other neuromodulator (Fig. 2b, 4e). This is consistent with previous reports that the neuropeptide POMC can be coexpressed with glutamate in some neurons, but GABA in others^32^^,^^33^. In our experiments, of five neurons expressing *Cck* (cholecystokinin), two coexpressed glutamate and one coexpressed GABA, and of twelve neurons expressing *Tac1* (tachykinin1), one coexpressed glutamate and two coexpressed GABA. We also found *Cck* and *Tac1* coexpressed with either glutamate or GABA in different neurons in another, large-scale study that analyzed single neurons from the hypothalamus^5^.

It is known that neurons expressing a neuropeptide or biogenic amine can also express a classical neurotransmitter ^4^^,^^5^^,^^31^^,^^34^^,^^35^. These results further indicate that many neuropeptides can be coexpressed with glutamate in some upstream neurons but with GABA in others. Of 34 neuropeptides coexpressed with glutamate or GABA in our studies, twelve were coexpressed with glutamate in some neurons and GABA in others (Fig. 4c).

These results raise the possibility that individual neuropeptides could exert different modulatory effects on CRHNs depending on whether they are released onto CRHNs together with glutamate or GABA. For example, if a neuropeptide acts to enhance CRHN excitability, it may further enhance glutamate stimulation of CRHNs, but dampen GABAergic suppression of CRHNs. Our observations suggest the possibility of a complex patterning of stimulatory, inhibitory, and neuromodulatory effects that act at the level of synaptic input to CRHNs to fine tune the effects of upstream inputs on the CRHNs.

### Upstream neurons can coexpress different combinations of neuropeptides

These studies also revealed many upstream neurons expressing different combinations of neuropeptides. As noted above, neuropeptides were expressed in 69% of the neurons upstream of CRHNs. While 38% of upstream neurons expressed a single neuropeptide, 32% coexpressed two or more, and a few neurons expressed 5-7 neuropeptides each (Fig. 5a, 5b). These results are consistent with previous reports that neurons expressing a given neuropeptide can also express another neuropeptide^36^^,^^37^, but the number of neuropeptides that could be coexpressed in single neurons was unexpected.

**Figure 5.**
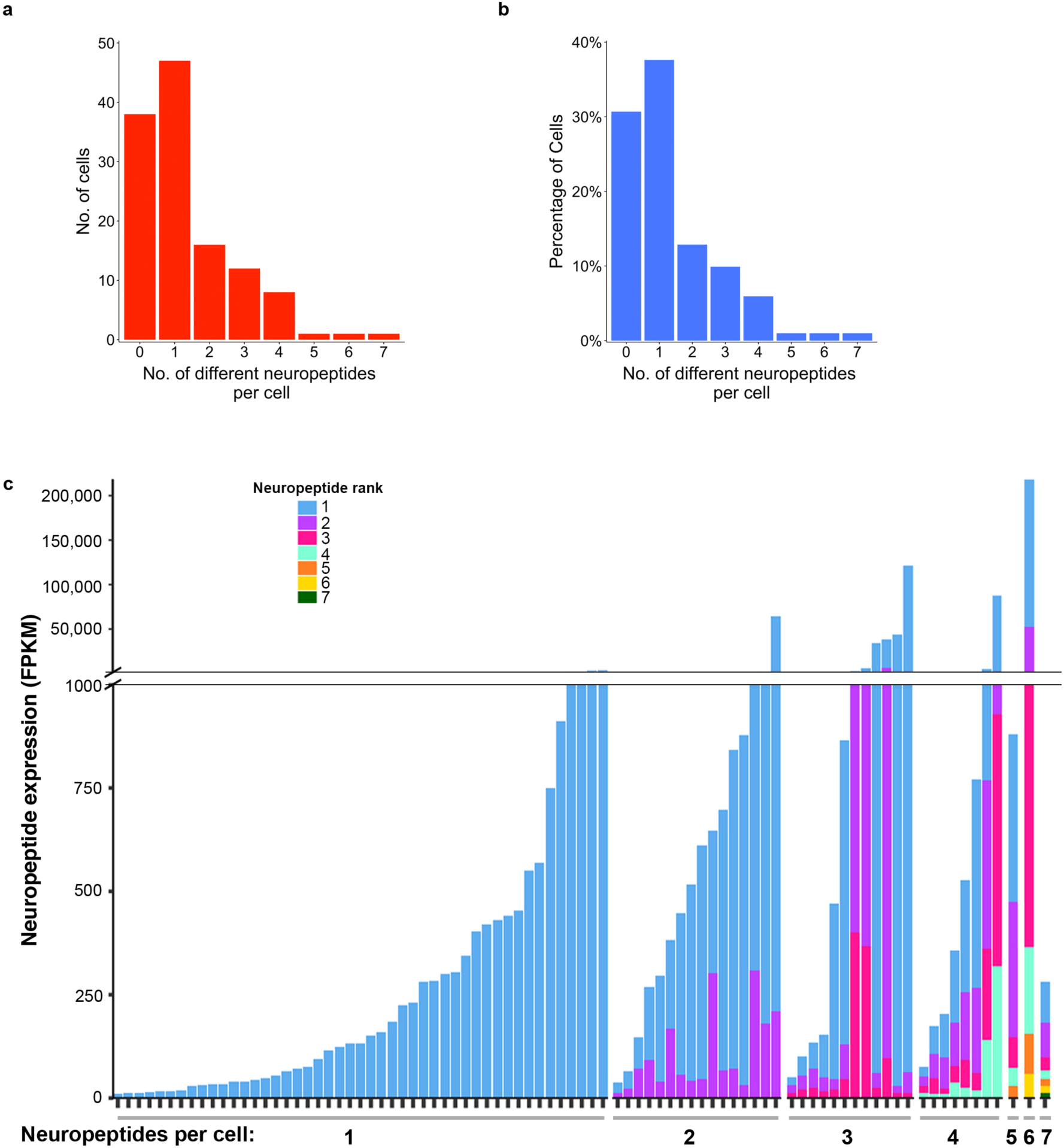
Upstream neurons can express different levels of more than one neuropeptide. **(a-b)** Graphs show the number (a) and percentage (b) of upstream neurons that expressed different numbers of neuropeptides. **(e**) Expression levels of neuropeptides in single neurons varied. Cells are arranged on the x-axis by the number of neuropeptides they expressed (1-7). Vertical bars indicate expression of neuropeptides in each cell, with different neuropeptides indicated in different colors and their expression in FPKM shown on the y-axis.

To further investigate the coexpression of neuropeptides in single neurons, we compared the levels of expression of different neuropeptides in individual neurons. At the level of single neurons, a few neurons expressed two or three neuropeptides at similar levels (Fig. 5c). However, many other neurons expressed one neuropeptide at a much higher level than one or more others expressed in the same cell. These results suggest that while a neuron might express multiple neuropeptides, the downstream effects of one might predominate over the effects of others. Some neuropeptides, as well as other signaling molecules, were expressed at very different levels in different cells (Fig. 6a, 6b), raising the possibility that they might play a more important role in signaling to CRHNs by some upstream neurons than others.

**Figure 6.**
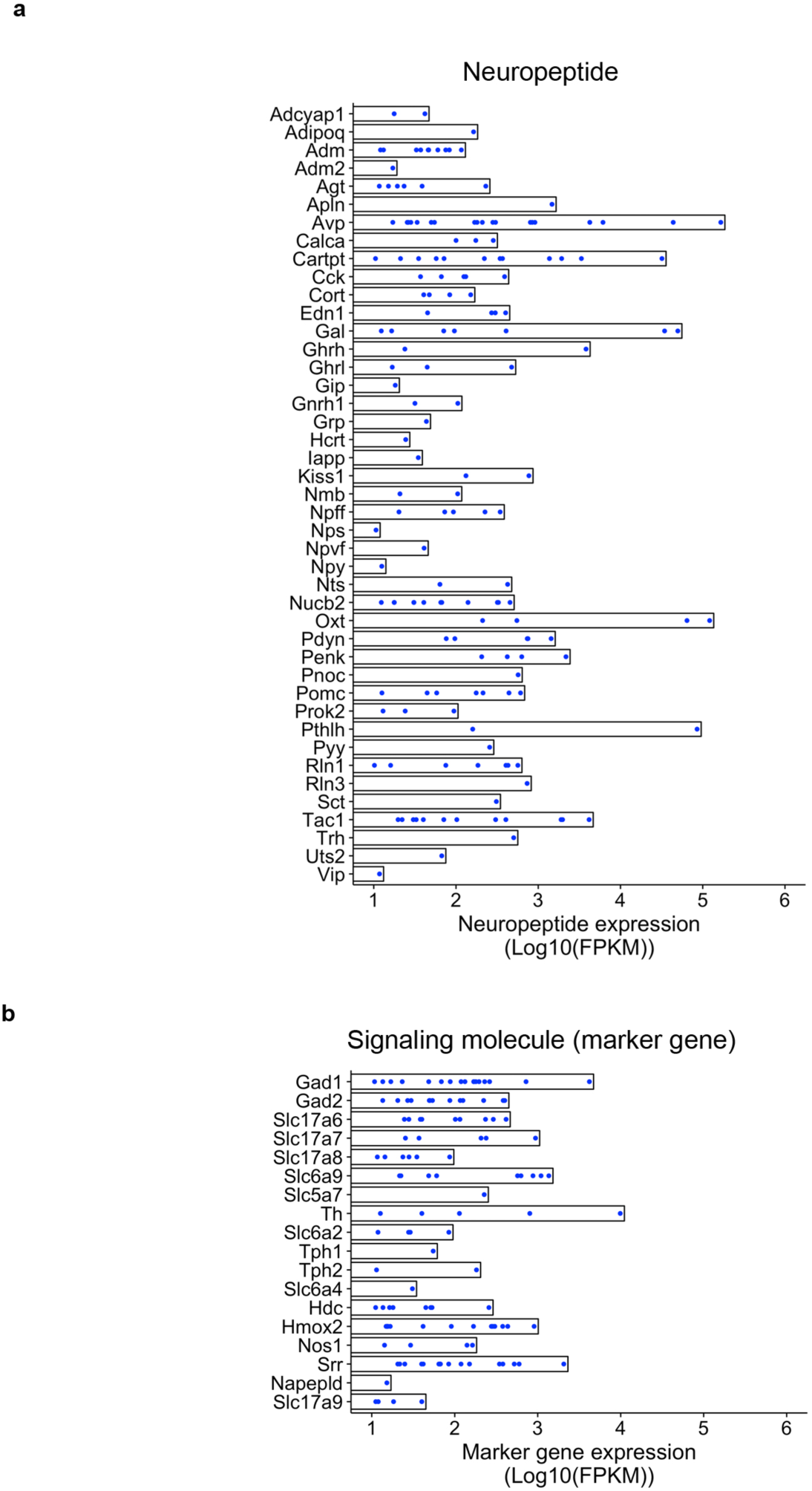
Expression levels of signaling molecules vary in upstream neurons. **(a-b)** Graphs show the expression levels of different neuropeptides (a) and marker genes for other signaling molecules (b) in individual neurons (blue dots) upstream of CRHNs. Data are shown as log-transformed FPKM.

Surprisingly, in addition to being expressed at different levels among neurons, neuropeptides were expressed in different combinations in different neurons (Fig. 7). For example, of seven neurons expressing *Gal* (galanin), *Avp* (arginine vasopressin) was coexpressed in five, *Cartpt* (cocaine- and amphetamine-regulated transcript protein) in three, and seven other neuropeptides each in 1-2 of the Gal+ neurons (Fig. 7a). In large-scale single cell RNA-seq data from the hypothalamus, we found that, altogether, 100 Gal-expressing neurons coexpressed nineteen other neuropeptides, including *Avp* and *Cartpt*^5^.

**Figure 7.**
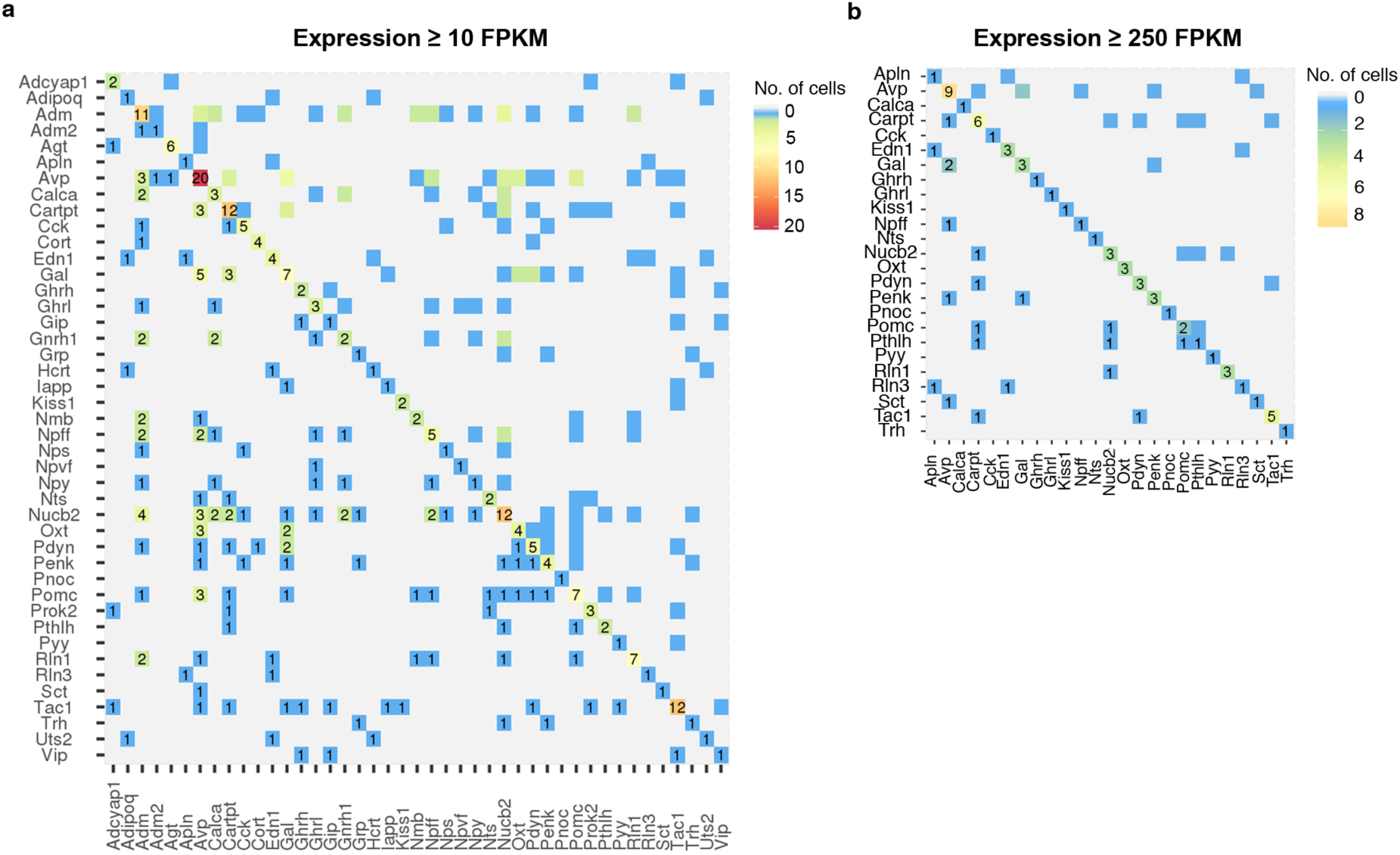
Upstream neurons can express diverse combinations of neuropeptides. **(a-b)** Coexpression of neuropeptides in upstream neurons detected at ≥10 FPKM (a) or ≥250 FPKM (b). Numbers and colored boxes indicate number of cells coexpressing a given pair of neuropeptides. Numbers on the diagonal show total cells expressing a given neuropeptide.

Using a higher transcript threshold to define gene expression in our experiments (≥250 FPKM (fragments per kilobase of transcript per million mapped reads) instead of ≥10 FPKM), the number of combinations of neuropeptides in different neurons was reduced, suggesting that much of the variability seen using the 10 FPKM threshold was the result of low level expression of neuropeptides (Fig. 7b). These results suggest that there may be more diversity in neuropeptide expression in single neurons than previously appreciated but that much of that diversity may result from the expression of neuropeptides at a low level.

### Upstream neurons expressing specific neuromodulators map to selected brain areas

One goal of these studies was to uncover molecular identifiers of upstream neurons that would allow for a molecular map to be superimposed on the anatomical map of neurons upstream of CRHNs. To investigate this possibility, we examined neurons upstream of CRHNs for the expression of several neuromodulators we had identified by transcriptome analysis of upstream neurons.

Using the Allen Brain Atlas (http://mouse.brain-map.org/)^38^, we examined the expression of a number of neuropeptides we had identified in different areas of the hypothalamus previously shown to contain neurons directly upstream of CRHNs^8^. Consistent with the fact that we had identified the neuropeptides in upstream neurons we isolated from the hypothalamus, all of the neuropeptides for which there was adequate information in the atlas were expressed in at least one of those hypothalamic areas in the atlas (Fig. 8a).

**Figure 8.**
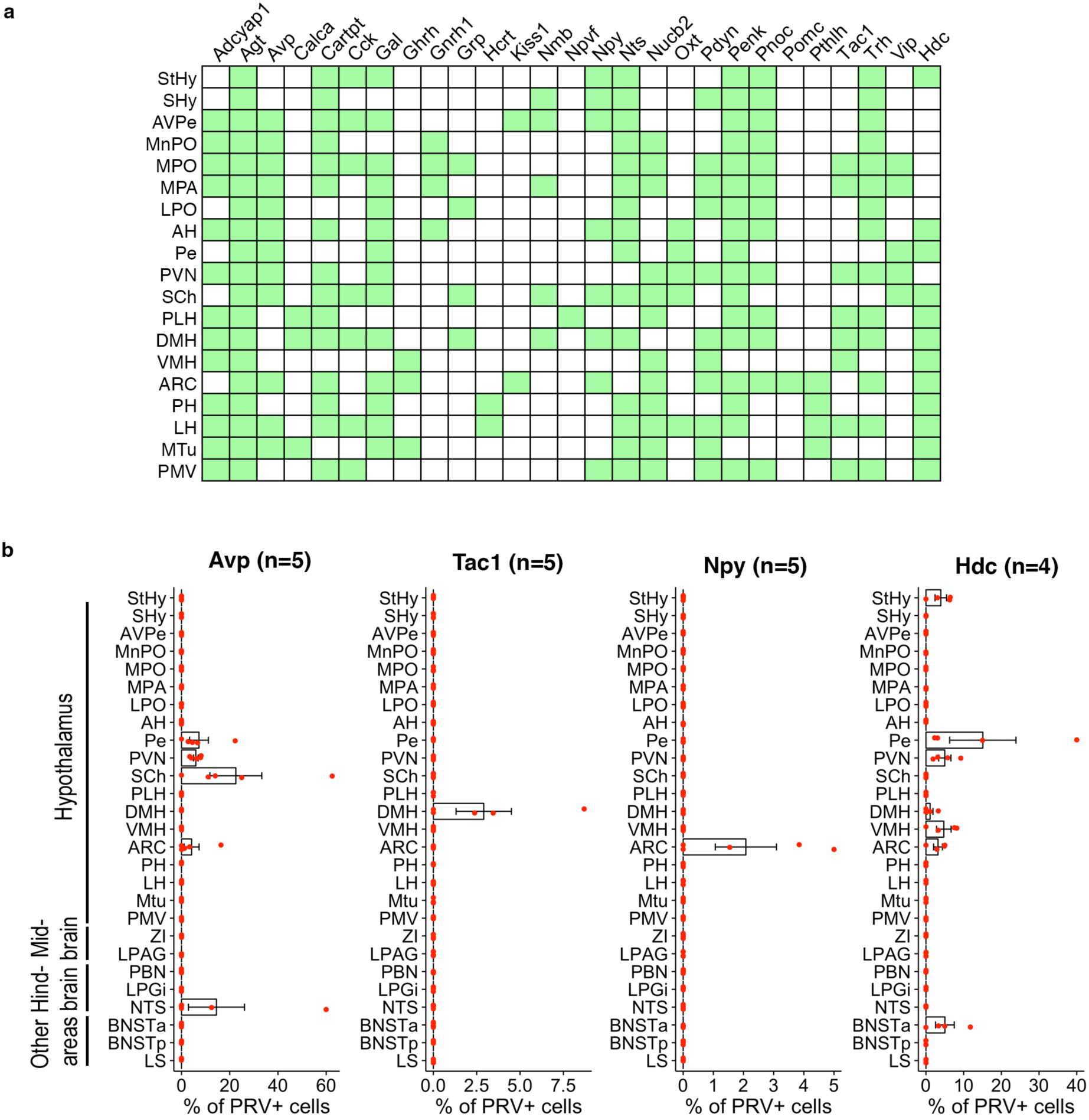
Upstream neurons expressing individual signaling molecules map to specific brain area(s). **(a)** Chart shows data obtained from the Allen Brain Atlas in situ hybridization database indicating expression of specific neuromodulators in areas of the hypothalamus that contain neurons upstream of CRHNs (green boxes). **(b)** Graphs show the percentage of PRV+ neurons colabeled for different neuromodulators (Avp, Tac1, Npy, or Hdc) in different brain areas following CRHN infection with PRVB177. Error bars indicate s.e.m.. Parenthesized numbers indicate the number of animals (n) per condition.

To analyze the locations of upstream neurons expressing specific neuromodulators, we first infected Cre-expressing CRHNs with PRVB177, which has Cre-dependent expression of hemagglutinin (HA)-tagged TK^8^. After 3 days, when the virus had crossed one synapse to infect immediately upstream neurons, we costained brain sections with riboprobes for specific neuromodulators and anti-HA antibodies to detect PRV-infected neurons (Fig. 8b and Supplementary Fig. 2).

*Avp* was detected in PRV+ neurons in five brain areas, including four areas of the hypothalamus (Fig. 8b). In contrast, *Tac1* (tachykinin 1) and *Npy* (neuropeptide Y) were each seen in PRV+ neurons in only one hypothalamic area, the DMH for *Tac1* and the ARC for *Npy*. *Hdc* (histidine decarboxylase), a marker for the expression of the biogenic amine, Histamine, was seen in six hypothalamic areas and one other brain area.

The selected expression of neuromodulators we identified in upstream neurons in only certain brain areas confirms that data obtained using Connect-seq can be used to superimpose a molecular map on the anatomical map of neural circuits upstream of CRHNs (Fig. 9). This information can be used to map functional responses to particular stressors to subsets of upstream neurons in specific brain areas, thereby providing insight into the functional meaning of the anatomical/molecular mapping and suggesting tools to dissect the roles of specific neuronal subsets in particular functions.

**Figure 9.**
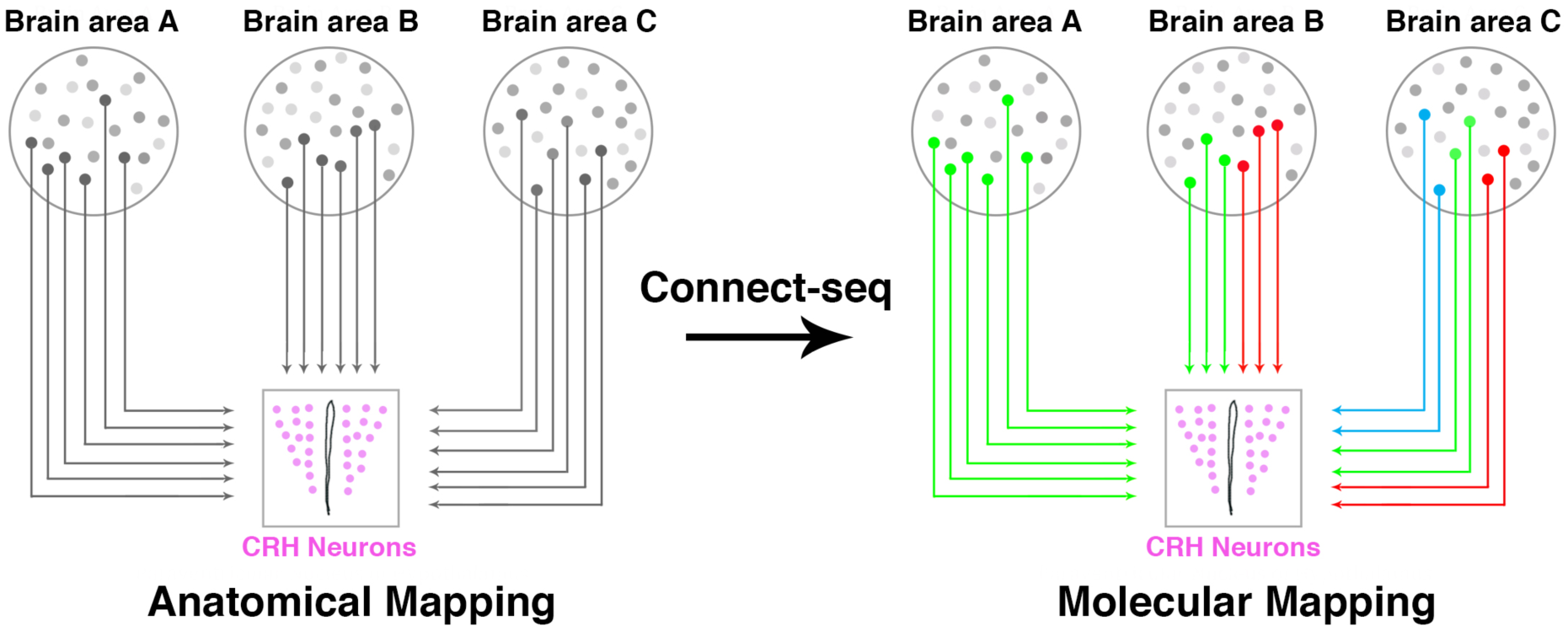
Connect-seq for superimposing molecular on anatomical circuit maps. Retrograde viral tracing provided an anatomical map of neurons upstream of CRHNs. Connectseq defined the transcriptomes of single neurons upstream of CRHNs and revealed signaling molecules they express. By mapping the locations of upstream neurons expressing those signaling molecules, it is possible to superimpose a molecular map on the anatomical map of neural circuits upstream of CRHNs.

## Discussion

The brain contains a vast number of neural circuits that govern diverse functions, but the neuronal components of those circuits and their individual contributions to circuit function are largely undefined^1^^-^^3^^,^^19^. In the present studies, we devised a strategy, termed ‘Connect-seq’, which utilizes a combination of viral tracing and single cell transcriptomics to uncover the molecular identities of neurons upstream of a specific set of neurons and reveal the neurotransmitters and neuromodulators they use to communicate with their downstream synaptic targets.

We used Connect-seq to gain insight into circuits that transmit information to CRH neurons that control blood levels of stress hormones. Neurons upstream of CRHNs expressed a remarkable number and variety of neurotransmitters and neuromodulators, including dozens of different neuropeptides. While some neurons expressed only one of these signaling molecules, others expressed different combinations, suggesting a complex array of input to CRHNs likely to excite or inhibit those neurons or modulate their excitability.

Localization experiments revealed subsets of upstream neurons expressing individual neuromodulators we had identified by Connect-seq in distinct brain areas. In conclusion, Connect-seq enables the construction of a molecular map that can be superimposed on an anatomical map of neural circuits, thereby allowing the investigation of roles played by individual neuronal components of those circuits under normal conditions and in disease.

## ACKNOWLEDGMENTS

We thank J. Delrow, A. Marty, A. Dawson, and R. Meredith at the Fred Hutchinson Cancer Research Center (FHCRC) Genomics Facility for their assistance with RNA-seq and A. Berger, S. Dozono, and B. Raden, at the FHCRC Flow Cytometry Facility for their assistance with flow sorting. We thank X. Ye, A. Spray, and E. Albrecht for technical assistance and members of the Buck Laboratory for helpful discussions. **Funding:** This work was supported by the Millen Literary Trust (E.J.L), the Howard Hughes Medical Institute (L.B.B), NIH grants R01 DC015032 (L.B.B), RO1 DC016442 (L.B.B.), and DP2 HD088158 (C.T), the Paul G. Allen Frontiers Foundation (through the Allen Discovery Center for Cell Lineage Tracing), and an Alfred P. Sloan Fellowship (C.T). L.B.B. is on the Board of Directors of International Flavors & Fragrances.

## Author contributions

N.K.H., K.K., and L.B.B. conceived the study; N.K.H. developed methods for isolation of PRV-infected cells and performed single-cell RNA-seq; E.J.L. performed PRV injections and staining for neuropeptides in upstream neurons; A.E. conducted computational analyses of RNA-seq data; K.K. designed and made PRVB180; D.K. provided riboprobes for experiments; R.B. aligned sequenced reads to the mouse and PRV genomes; N.K.H., E.J.L., A.E., and L.B.B. analyzed the data; and N.K.H. and L.B.B. wrote the paper. The supplementary information contains additional data.

## Methods

### Mice

CRH-IRES-Cre (CRH-Cre) mice were generated as described previously^13^. All procedures involving mice were approved by the Fred Hutchinson Cancer Research Center Institutional Animal Care and Use Committee. C57BL/6J mice were purchased from The Jackson Laboratory.

### Pseudorabies Viral Vectors

Construction of PRVB177 was described previously^8^. PRVB180 construct was made according to the methods described previously^8^. Briefly, PRVB180 was constructed using homologous recombination between targeting vector (PRVTK-GFP) and genomic DNA of PRV TK-BaBlu, a thymidine kinase (TK)-deleted PRV Bartha strain derivative with a LacZ insertion into the gG locus^39^. For generating PRVTK-GFP, a flexstop-flanked sequence^40^ encoding a PRV TK fused at its C terminus to enhanced green fluorescent protein (eGFP), obtained from pEGFP-N1 vector (Clontech), was first cloned with an inverse orientation into an eGFP-deleted pEGFP-N1 vector (Clontech). Next, following restriction digestion, NsiI fragments containing a CMV promoter, the flexstop-flanked coding sequence, and an SV40 polyadenylation signal were cloned between gG locus sequences matching those 5′ and 3′ to the lacZ sequence in PRV TK-BaBlu to give the final targeting vector (PRVTK-GFP). These vectors were then linearized and co-transfected with PRV TK-BaBlu genomic DNA into HEK 293T cells (ATCC). Recombinant virus clones were selected and confirmed following methods described previously^41^.

PRVs were propagated by infecting PK15 cells (ATCC) with the viruses using a multiplicity of infection (m.o.i.) = 0.1– 0.01. After ~2 days of infection, cells showed a prominent cytopathic effect. Cells were scraped from the dishes, pelleted by centrifugation, and frozen using liquid nitrogen and then quickly thawed in a 37°C water bath. After three freeze-thaw cycles, cell debris was removed by centrifugation twice at 1,000g for 5 min, and the supernatant was aliquoted and stored at -80°C, until use. The titer of viral stocks was determined using standard plaque assays on PK15 cells^42^ with titers expressed in plaque-forming units (p.f.u.).

### Stereotaxic injections

Viruses were injected into the PVN of CRH-Cre mice as described previously^8^. All injections were done under inhalation anesthesia of 2% isoflurane. Briefly, 1ul of PRVs (PRVB180 & PRVB177; 1–1.5 × 10^6^ p.f.u.) were loaded into a 1ul syringe and were injected into the brain at a rate of 100nl per minute using a Stereotaxic Alignment System (David Kopf Instruments). The needle was inserted to the PVN based on a stereotaxic atlas^43^. After recovery, animals were singly housed with regular 12 h dark/light cycles, and food and water were provided ad libitum.

### Isolation of PRV-infected single cells using flow cytometry

Adult CRH-Cre mice aged 2-4 month were injected with PRVB180. After three days, mice were euthanized by cervical dislocation, the brain quickly removed and submerged into ice-cold Hibernate-A medium (#A1247501; ThermoFisher Scientific). The hypothalamus was carefully microdissected under microscope and dissociated into tiny pieces in dissociation buffer containing Hibernate-A medium with papain (10 U ml^-1^, PAP2, PDS kit; Worthington Biochemical Corporation) and DNase (200 U ml^-1^, DNase vial, D2, PDS kit; Worthington Biochemical Corporation)). Dissociated hypothalamic tissue were transferred into 5 ml of dissociation buffer and incubated for 30 min at room temperature. After incubation, tissue pieces were gently triturated 2-3 times through a series of fire-polished glass Pasteur pipettes with decreasing diameters of ~600 µm, ~300 µm, and ~150 µm to dissociate tissue into a cloudy suspension. Cells were sieved using a 70 µm cell strainer into 50 ml tubes containing 10 ml of ice-cold Hibernate-A medium with 1% BSA. Cells were then centrifuged at 800 rpm for 5 min at 4 °C, washed once with ice-cold Hibernate-A medium and cell pellet obtained was resuspended in ~500 µl of Hibernate-A medium containing 2% B27 (ThermoFisher) and DAPI (0.5 ng ml^-1^) to stain for dead cells. Cells were then sieved using a 40 µM cell strainer to obtain single cell suspension and proceeded to FACS (fluorescence-activated cell sorting).

PRV-infected single cells were isolated based on the fluorescence emitted by TK-GFP using flow cytometry (FACSAria II; BD Biosciences) according to methods described previously^14^ with minor modifications. Briefly, cells were first sorted based on their size to isolate single cells using Forward (FSC) and Side Scatter (SSC) gates, and then high GFP and low DAPI (to obtain live cells) cells were individually placed in 96 well plates containing 3 µl of RLT Plus lysis buffer (#1053393; Qiagen).

### Single-cell RNA Sequencing

Single cells were then processed immediately to generate cDNA libraries using SMART-seq2^15^^,^^16^ with minor modifications. Briefly, polyadenylated mRNAs were captured by incubating cell lysates for 20 min at room temperature with a modified biotinylated oligo-dT primer (5’-biotin-triethyleneglycol-AGCAGTGGTATCAACGCAGAGTACT30VN-3’, where V is either A, G, or C, and N is any base, HPLC purified; Eurofins Genomics LLC) conjugated to Streptavidin-coupled magnetic beads (#11205D, Dynabeads; ThermoFisher) for 20 min at room temperature. Biotinylated oligo-dT primer was conjugated to Streptavidin-coupled magnetic beads according to the manufacturer’s instructions. After incubation, the plates were placed on a magnet, retaining the beads with mRNAs at the bottom, washed twice with wash buffer containing 1X Superscript II first strand buffer, 2 U µl^-1^ RNAse inhibitor (#30281, Lucigen) and 0.5% (vol/vol) Tween 20 (#P9416; Sigma-Aldrich) in Nuclease-free water (#AM997; ThermoFisher) at room temperature and resuspended in 11 µl of reverse transcription mix containing 1X Superscript II first-strand buffer, 1 mM dNTP (#R1092; ThermoFisher), 6 mM MgCl_2_, 1M Betaine (#B0300; Sigma-Aldrich), 5 mM DTT, 1 µM template-switching oligo (TSO, AAGCAGTGGTATCAACGCAGAGTGAATrGrG+G, RNase-free HPLC purified; Exiqon), 0.5 U µl^-1^ RNAse inhibitor and 10 U µl^-1^ Superscript II reverse transcriptase (#18064-014, ThermoFisher) in Nuclease-free water. mRNAs were then reverse transcribed by incubation at 42°C for 90 min followed by 10 cycles of 50°C for 2 min, 42°C for 2 min, and then incubated at 70°C for 15 min. The resulting cDNAs were then amplified using 1X KAPA HiFi HotStart ReadyMiX (#KK2602, KAPA Biosystems) and IS PCR primer (AAGCAGTGGTATCAACGCAGAGT, HPLC purified; Eurofins Genomics LLC) for 30 PCR cycles.

Amplified cDNAs were then purified and size selected using equal volume of AMPure XP beads (#A63880; Beckman Coulter). cDNA libraries were verified for the presence of PRV transcripts by PCR using primers for GFP (5’, GAGCAAGGCGAGGAGCTGTT, 3’ GGTCAGCTTGCCGTAGGTG). Purified cDNA libraries were then quantified using the Qubit dsDNA HS assay kit (#Q32854; ThermoFisher Scientific). Sequencing libraries were then generated using ¼ reaction volume of the Nextera XT DNA library prep kit (Illumina) according to manufacturer’s instructions. Briefly, 1.25 µl of single-cell cDNAs (~0.25 ng µl^-1^) were used for tagmentation reaction, and then PCR amplified to insert sequencing adaptors and cell-specific barcodes.

The resulting libraries were then subjected to Illumina deep sequencing. Sequencing was performed using an Illumina HiSeq 2500 instrument in rapid mode employing a paired-end, 50 base read length (PE50) sequencing strategy. Image analysis and base calling were performed using Illumina Real Time Analysis v1.18 software, followed by “demultiplexing” of indexed reads and generation of FASTQ files using Illumina’s bcl2fastq Conversion Software (v1.8.4). Reads of low quality were discarded prior to adapter trimming using Trim Galore (v.0.4.4, available at **http://www.bioinformatics.babraham.ac.uk/projects/trim_galore/**) with the options “-- adapter AAGCAGTGGTATCAACGCAGAGTAC --stringency 8 --quality 0 -e 0.15 --length 20 -- paired --retain_unpaired”.

The default options in TopHat^17^ (v2.1.0) were used to align reads to the mouse genome (UCSC mm10 assembly, using GENCODE M15 release gene models) as well as to the Pseudorabies virus genome (NC_006151.1). Cufflinks^18^ (v.2.2.1) was used to estimate gene expression profiles in units of fragments per kilobase of transcript per million mapped reads (FPKM), by first running the “cuffquant” tool on the aligned reads for each cell with the “-u” option, which performs additional algorithmic steps designed to better assign ambiguously mapped reads to the correct gene of origin. Per-cell gene expression profiles were subsequently normalized with the “cuffnorm” utility, using the “classic-fpkm” normalization method, for use in downstream analysis. A total of 384 PRV-infected cells were sequenced at a depth of at least 7,000,000 reads (median ~ 6,700,000, range ~17,500 to ~13,300,000). 347 cells that expressed at least 500 genes per cell were considered for downstream analysis. On average, cells expressed ~2,600 genes per cell (median ~2,500, range ~520 to 10,300).

### Analysis of marker genes

Transcriptome data were manually inspected for transcripts of cell type markers to include neurons and exclude non-neuronal cells. We used a threshold of ≥10 FPKM for cell type markers as in our previous analysis of olfactory sensory neurons^44^. Cells were classified using a criterion that a cell must express two or more of six different marker genes of a cell type but not more than two of other cell types. We used the following genes to classify cells: Snap25, Map2, Syp, Kif5c, Dlg2, and Dlg4 for neurons; Aldh1l1, Aldoc, Aqp4, Gfap, S100b, and Sparcl1 for astrocytes; Adgre1, Aif1, Tmem119, Cxcr1, Hexb, and Mafb for microglia; Cldn11, Mbp, Mobp, Mog, Pdgfra, and Gpr17 for oligodendrocytes. If a cell remained unclassified but expressed one marker of neurons but not other cell types, we additionally examined markers for the excitatory neurotransmitter glutamate (Slc17a/6/7/8) and the inhibitory neurotransmitter GABA (Gad1/2) and included cells with one or more of those markers. For neurotransmitters and neuromodulators, we used marker genes that encode rate-limiting enzymes required for their biosynthesis or vesicular transporters that package synaptic vesicles with specific neurotransmitters. For neuropeptides, we used genes encoding known ligands of GPCRs obtained from the International Union of Basic and Clinical Pharmacology (IUPHAR)/ British Pharmacological Society (BPS) website (http://www.guidetopharmacology.org) and those with evidence for having an electrophysiological effect on neurons.

### In situ hybridization

Dual labeling studies to examine the expression of markers in PRV-infected cells were performed essentially as described previously^8^^,^^44^. Coding region fragments of Avp, Tac1, Npy, and Hdc were PCR-amplified from mouse brain cDNA or genomic DNA and cloned into the pCR4 Topo vector (Thermo Fisher). Digoxigenin (DIG)- labeled riboprobes were prepared using the DIG RNA Labeling Mix (Roche). The PVN of CRH-Cre mice aged 2-4 months was injected with PRVB177, as described above. After three days, mice were euthanized by cervical dislocation, and brains were frozen in OCT (Sakura) on dry ice and stored at -80 °C. Coronal cryostat sections of 20µm were hybridized to DIG-labeled riboprobes at 56 °C for 13–16 h.

#### Costaining for HA (PRVB177) and neuropeptides riboprobes

After hybridization with riboprobes, sections were washed twice for 5 min at 63 °C in 5x SSC followed by twice for 30 min at 63 °C in 0.2x SSC. Sections were then incubated with horseradish peroxidase (POD)-conjugated sheep anti-DIG antibodies (1:200; 1207733910; Roche) and biotinylated anti-HA antibodies (1:300, #901505; BioLegend) diluted in blocking buffer (1% Blocking reagent, #FP1012; Perkin Elmer) for 2h at 37 °C. Sections were then washed three times for 5 min at RT in TNT (0.1M Tris-HCl pH7.5, 0.5M NaCl, 0.05 % Tween) buffer, and then treated using the TSA-plus FLU kit (Perkin Elmer). Sections were then incubated with 0.5 µg ml^−1^ DAPI and Alexa555-Streptavidin (1:1,000, #32355; ThermoFisher) at room temperature for 1 h, and then coverslipped with Fluoromount-G (#0100-01; Southern Biotech)

### Cell counting

Sections were analyzed essentially as described previously^8^. Briefly, images were collected using an AxioCam camera and AxioImager.Z2 microscope with an apotome device (Zeiss). A mouse brain atlas was used to identify brain structures microscopically and in digital photos. Every fifth section was analyzed for all experiments.

Abbreviations for brain areas. Abbreviations used for brain areas are according to a mouse brain atlas^43^ and our previous report^8^. AH, anterior hypothalamic area; ARC, arcuate hypothalamic nucleus; AVPe, anteroventral periventricular nucleus; BNSTa, bed nucleus of the stria terminalis, anterior part; BNSTp, bed nucleus of the stria terminalis, posterior part; DMH, dorsomedial hypothalamic nucleus; LH, lateral hypothalamic area; LPAG, lateral periaqueductal grey; LPGi, lateral paragigantocellular nucleus; LPO, lateral preoptic area; LS, lateral septal nucleus; MnPO, median preoptic nucleus; MPA, medial preoptic area; MPO, medial preoptic nucleus; MTu, medial tuberal nucleus; NTS, nucleus of the solitary tract; PBN, parabrachial nucleus; Pe, periventricular nucleus of the hypothalamus; PH, posterior hypothalamic nucleus; PLH, peduncular part of lateral hypothalamus; PMV, premammillary nucleus, ventral part; SCh, suprachiasmatic nucleus; SHy, septohippocampal nucleus; StHy, striohypothalamic nucleus; VMH, ventromedial hypothalamic nucleus; ZI, zona incerta.

---

**Supplementary Figure 1.**
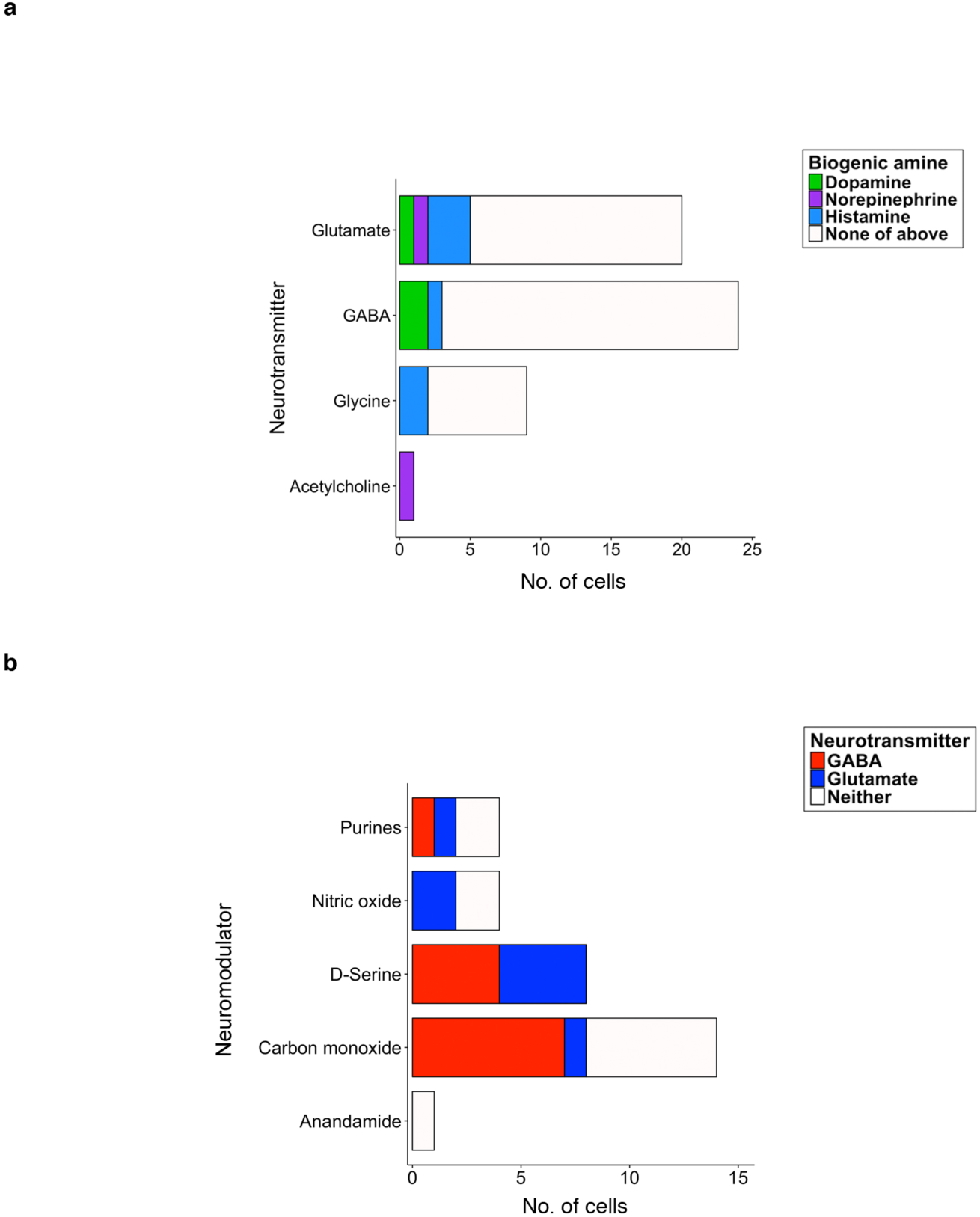
Coexpression of different signaling molecules in upstream neurons. Graphs show the number of upstream neurons that coexpressed different neurotransmitters with different biogenic amines (a) or that coexpressed other neuromodulators with GABA or glutamate (b).

**Supplementary Figure 2.**
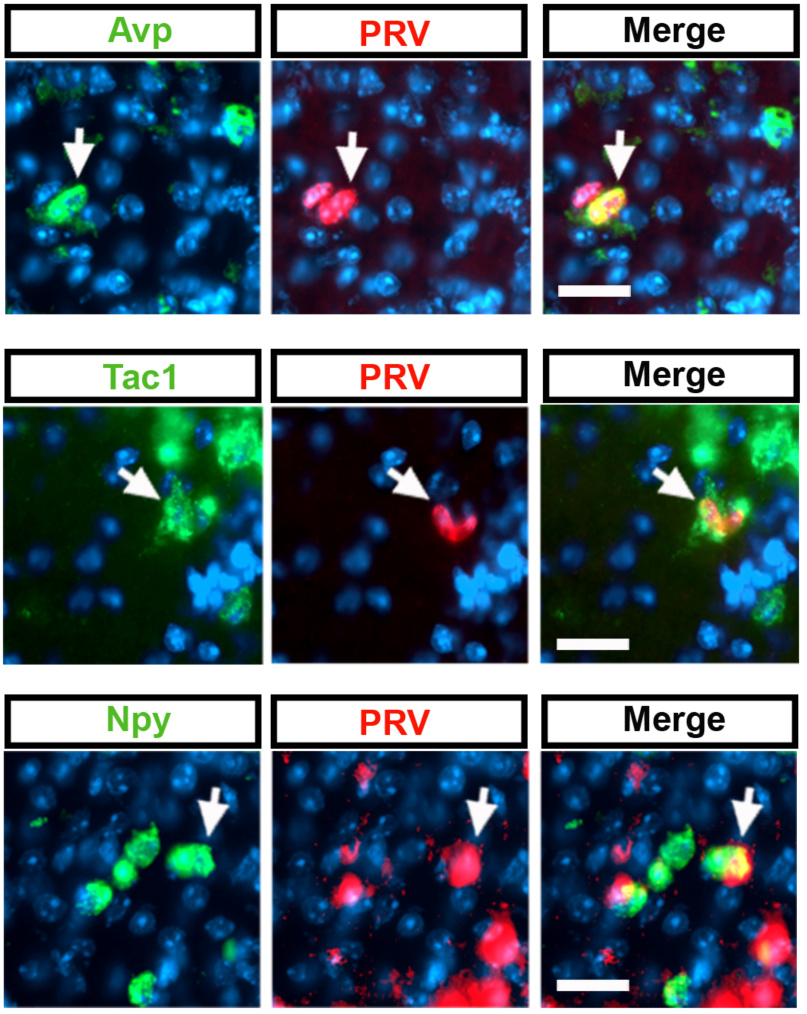
Expression of genes encoding neuropeptides in neurons upstream neurons of CRHNs. CRHNs were infected with PRVB177 and brain sections costained on d3pi with neuropeptide riboprobes (green) and anti-HA antibodies (red) (PRV+ cells). Arrows indicate colabeled neurons. Scale bars, 25 µM.

